# Chitosan induces plant hormones and defences in tomato root exudates

**DOI:** 10.1101/2020.06.09.142653

**Authors:** Marta Suarez-Fernandez, Frutos Carlos Marhuenda-Egea, Federico Lopez-Moya, Marino B. Arnao, Francisca Cabrera-Escribano, Maria Jose Nueda, Benet Gunsé, Luis Vicente Lopez-Llorca

## Abstract

In this work, we use electrophysiological and metabolomic tools to determine the role of chitosan as plant defence elicitor in soil for preventing or manage root pests and diseases sustainably. Root exudates include a wide variety of molecules that plants and root microbiota use to communicate in the rhizosphere. Tomato plants were treated with chitosan. Root exudates from plants were analysed at 3, 10, 20 and 30 days after planting (dap). We found, using High Performance Liquid Chromatography (HPLC) and Excitation Emission Matrix (EEM) fluorescence, that chitosan induces plant hormones, lipid signalling and defence compounds in tomato root exudates, including phenolics. High doses of chitosan induce membrane depolarization and affect membrane integrity. ^1^H-NMR showed the dynamic of exudation, detecting the largest number of signals in 20 dap root exudates. Root exudates from plants irrigated with chitosan inhibit ca. 2-fold growth kinetics of the tomato root parasitic fungus *Fusarium oxysporum* f. sp. radicis-lycopersici. and reduced ca. 1.5-fold egg hatching of the root-knot nematode *Meloidogyne javanica*.

**One-sentence summary:** Chitosan depolarizes plasma membrane of root cells, causing the secretion of hormones, lipid signalling and plant defence compounds, including phenolics. These root exudates inhibit soil-borne pathogens.

## INTRODUCTION

Chitosan is a linear polymer of β-(1-4)-linked N-acetyl-2-amino-2-deoxy-D-glucose (acetylated) and 2-amino-2-deoxy-D-glucose (deacetylated) (Kaur and Dhillon, 2014). It is generally obtained by partial deacetylation of chitin (Ravi Kumar, 2000), which is the second most abundant polysaccharide in nature after cellulose (Elieh-Ali-Komi and Hamblin, 2016). Chitin is a major component of the cuticle of insects, exoskeleton of crustaceans and fungal cell walls. Chitosan has been described as elicitor of plant defences (Yin *et al.*, 2016) and it is used in food crops such as tomato and tobacco (Iriti and Faoro, 2008; El-Tantawy, 2009). Chitosan also propitiates accumulation of auxin (mainly indoleacetic acid; IAA) in the apex of plant roots (Lopez-Moya *et al.*, 2017). Electrophysiology can monitor the response of plant roots to stress (Rodrigo-Moreno *et al*., 2013). Membrane potential reflects the action of all pumps in the cell membrane to maintain ion gradients (Alberts *et al.*, 2002). Root cell membranes detect changes in their environment and respond starting metabolic cascade reactions (Fürstenberg-Hägg *et al*., 2013; Matzke and Matzke, 2013). Those reactions could lead to new compounds of agronomic and ecological interest.

Rhizodeposition is the process of releasing organic compounds from roots to the external medium. Plants exude a wide variety of low molecular weight organic compounds (e.g. amino acids and small peptides, organic acids, hormones, sugars, phenolics and other secondary metabolites) (Vivanco *et al*., 2002). Resolving this complex mixture requires the use of diverse metabolomics technologies (Escudero *et al.*, 2014; van Dam and Bouwmeester, 2016). Root exudates are paramount in plant-microbe interactions in the rhizosphere, including beneficial and pathogenic microbes and play a key role in signalling (Hirsch *et al.*, 2003; Bais *et al.*, 2006; Walker *et al*., 2009; Zhang *et al*., 2019; Valette *et al.*, 2020). Metabolomics can help us to understand the chemical interactions between organisms in the rhizosphere, as well as the importance to uncover toxic compounds (Baldrian, 2019). Metabolomics allows detection of phenolic compounds by fluorescence (Hupp *et al.*, 2019). Phenolics have diverse functions in plants, but one of the most important is their role in plant defence and signalling (Mandal *et al.*, 2010). Hormones are also signalling molecules that can also be found using metabolomics (Street and Schenk, 1981; Li *et al.*, 2009; Yang *et al*., 2015). Auxin (e.g. IAA), salicylic acid (SA), jasmonic acid (JA) and abscisic acid (ABA) are produced in response to physiologic or metabolic changes (Asami and Nakagawa, 2018). Phytomelatonin is considered a master regulator in plant stress conditions (Arnao and Hernández-Ruiz, 2019a; Arnao and Hernández-Ruiz, 2019b; Arnao and Hernández-Ruiz, 2019c) and also involved in the regulation of several plant hormones (Arnao and Hernández-Ruiz, 2018). Phytomelatonin promotes growth and root appearance (Hernández-Ruiz *et al.*, 2005; Arnao and Hernández-Ruiz, 2007).

The aim of this work is to determine the effect of chitosan on plant rhizodeposition. Root electrophysiology allows us to monitor the effect of chitosan on membrane functionality. In addition, metabolomic techniques are used to determine the effect of how chitosan modulates the composition of root exudates. Finally, we test these exudates against root pathogens which threaten food security worldwide. This will allow us to validate the role of chitosan as a plant defence inducer in soil for preventing or managing root pests and diseases sustainably.

## RESULTS

### Chitosan depolarizes plasma membrane and reduces tomato root cell viability

Chitosan (1 and 2 mg·mL^-1^) depolarizes (p<0.001) plasma membrane of tomato root cells (Figure 1A). This is reflected in loss of cell viability (PI red staining, Figure 1B). Roots incubated with 2 mg·mL^-1^ chitosan show a curved morphology and dark precipitates. Conversely, a low dose of chitosan (0.1 mg·mL^-1^) does not alter plasma membrane potential. However, these roots show both viable (FDA green staining) and non-viable cells, indicating some chitosan damage. Untreated tomato roots show mostly viable cells, stained with FDA. The slight red staining is due to natural senescence of root epidermic cells (Figure 1B, arrow).

**Figure 1.**
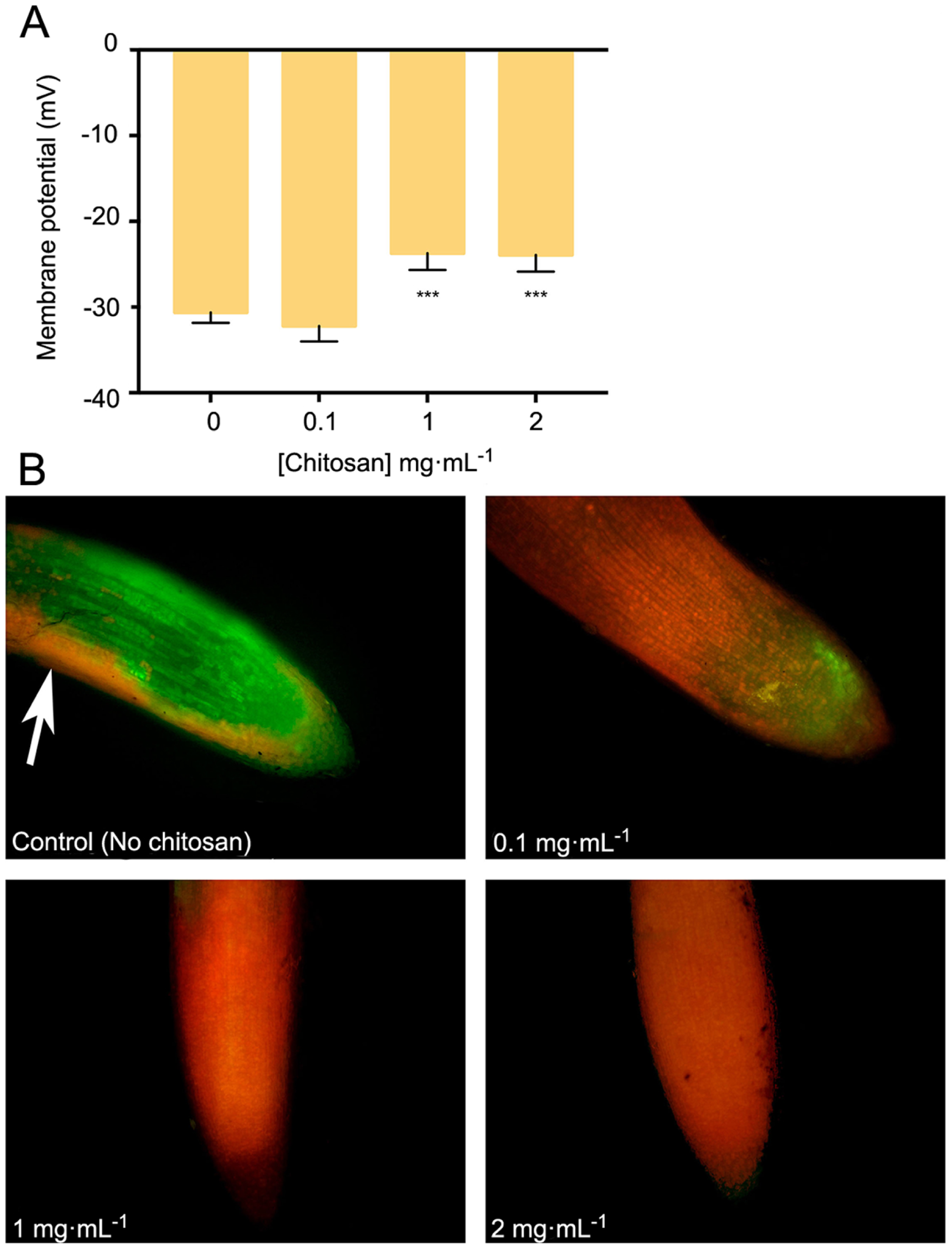
Chitosan depolarizes plasma membrane and damages tomato root cells. A, Variation in membrane potential of root cells with chitosan. High doses of chitosan (2 mg·mL^-1^) significantly reduce membrane potential. B, High doses of chitosan damage root cells after 24 h. Red staining labels damaged cells while green staining labels living ones. Multifactorial ANOVA was used to compare treatments (p-values 0.05 (*), 0.01 (**), 0.001 (***) and 0.0001 (****)).

### Chitosan induces hormones and phenolic compounds in root exudates

EEM Fluorescence analysis of root exudates resulted in 3 components (Figure 2, Supplemental Table 1). Components 1 and 3 are the most induced (p<0.01) by chitosan. Component 1 includes a putative fluorophore with Ex/Em wavelength pair of 315/430 nm, which could correspond to SA (Street and Schenk, 1981). Component 3 includes Ex/Em wavelength pairs of 245/384 and 265/384 nm, which may correspond to aromatic amino acids and peptides (Yang *et al.*, 2015; Table 1).

**Table 1.** Excitation and emission pairs of coordinates for each PARAFAC model per time. Abbreviations SA = Salicylic Acid, IAA = Indole Acetic Acid, dap = days after planting, C1 = Component 1, C2 = Component 2, C3 = Component 3.

| Time | Excitation (nm) | Emission (nm) | Compound |
| --- | --- | --- | --- |
| 3 dap | C1: 240/315<br>C2: 230/280<br>C3: 245/265/280 | C1: 430<br>C2: 336<br>C3: 384 | C1: SA (Street and Schenk, 1981)<br>C2: Phenolics (Mostofa *et al.*, 2013; Parri *et al.*, 2020)<br>C3: Aromatic amino acids and peptides (Yang *et al.*, 2015) |
| 10 dap | C1: 240/310<br>C2: 285<br>C3: 230/275 | C1: 436<br>C2: 364<br>C3: 312/358 | C1: SA (Street and Schenk, 1981)<br>C2: IAA (Li, 2009)<br>C3: Aromatic amino acids and peptides (Yang *et al.*, 2015), Phenolics (Mostofa *et al.*, 2013; Parri *et al.*, 2020) |
| 20 dap | C1: 240/310<br>C2: 285<br>C3: 230/270 | C1: 438<br>C2: 366<br>C3: 308 | C1: SA (Street and Schenk, 1981)<br>C2: IAA (Li, 2009)<br>C3: Phenolics (Mostofa *et al.*, 2013; Parri *et al.*, 2020) |
| 30 dap | C1: 235/310<br>C2: 230/280 | C1: 440<br>C2: 332 | C1: SA (Street and Schenk, 1981)<br>C2: Aromatic amino acids and peptides (Yang *et al.*, 2015), Phenolics (Mostofa *et al.*, 2013; Parri *et al.*, 2020) |

**Figure 2.**
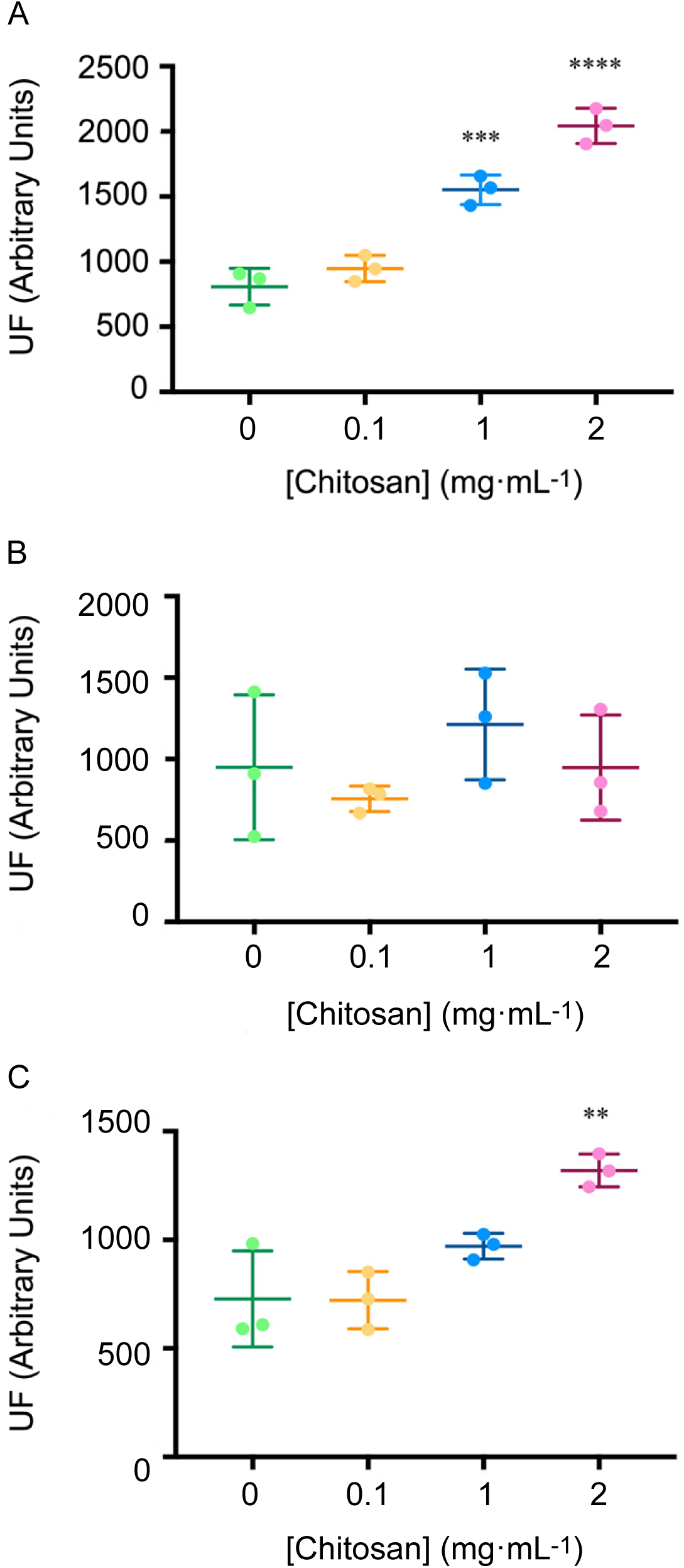
Chitosan increases EEM fluorescence of tomato root exudates. A, Component 1 (Salicylic Acid); B, Component 2 (Phenolics and Salicylic Acid derivatives); C, Component 3 (Aromatic aa and Peptides). For Ex/Em coordinates of Components see Table 1. Abbreviations: UF = fluorescence units, EEM = Emission Excitation Matrix. Multifactorial ANOVA was used to compare treatments (p-values 0.05 (*), 0.01 (**), 0.001(***) and 0.0001(****)).

Chitosan induces other hormones in tomato root exudates (Figure 3). High doses of chitosan (1 and 2 mg·mL^-1^) induce (p<0.01) IAA accumulation in root exudates (Figure 3A). Plant defence hormones (SA, JA and ABA) are also significantly induced (p<0.05) by chitosan (1 mg·mL^-1^) in tomato root exudates (Figures 3B-3D). This effect is lost at 2 mg·mL^-1^ chitosan. This could be the result of a root systemic damage caused by large chitosan concentrations. In view of the effect of chitosan on plant hormone homeostasis, endogenous phytomelatonin in roots was evaluated. Chitosan (1 mg·mL^-1^) causes a slight rise on phytomelatonin content in tomato root cells (Figure 3E). Phytomelatonin levels detected in tomato root exudates are close to nil (Supplemental Figure 1).

**Figure 3.**
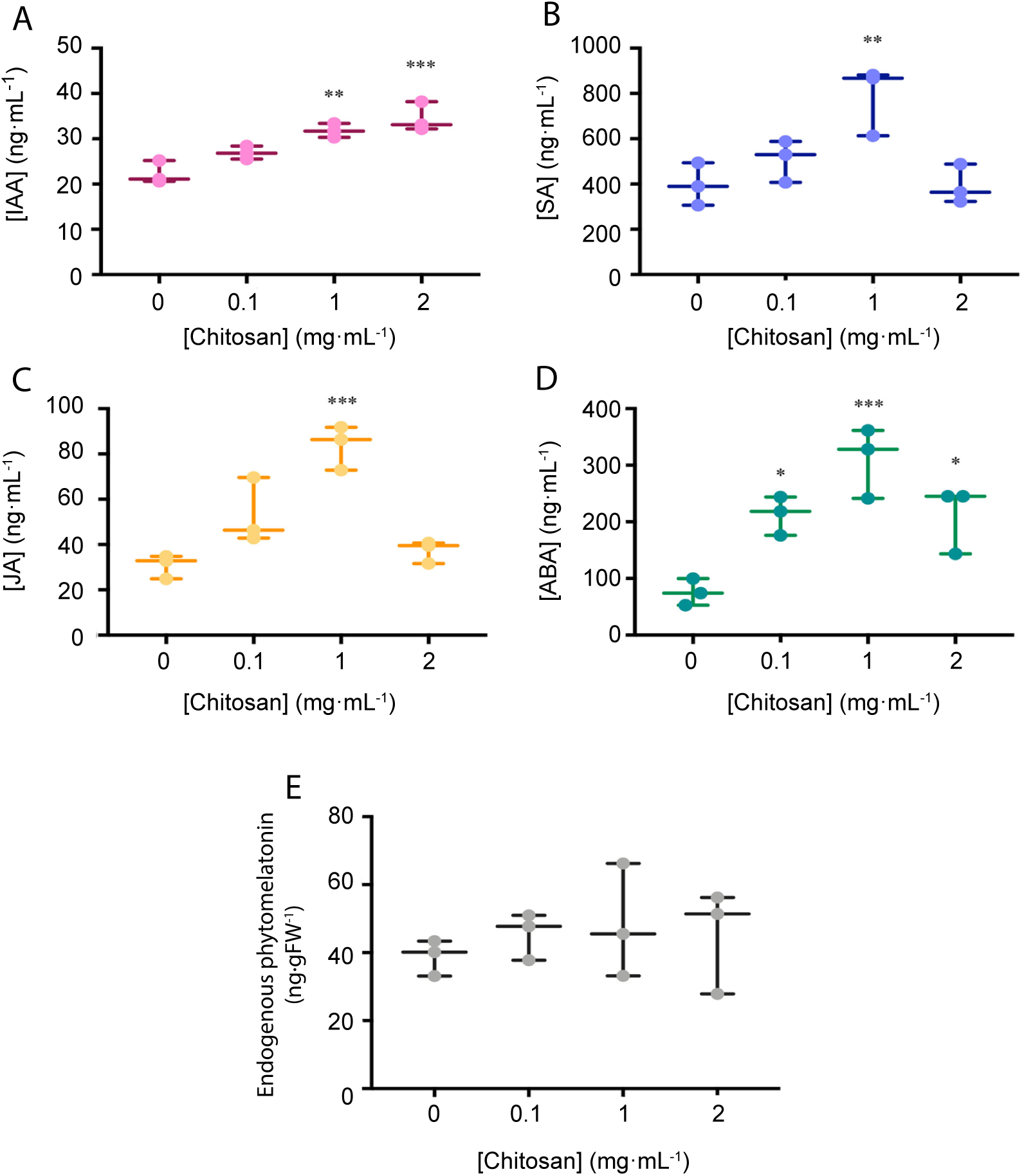
Chitosan increases EEM fluorescence of tomato root exudates. A, Component 1 (Salicylic Acid); B, Component 2 (Phenolics and Salicylic Acid derivatives); C, Component 3 (Aromatic aa and Peptides). For Ex/Em coordinates of Components see Table 1. Abbreviations: UF = fluorescence units, EEM = Emission Excitation Matrix.-Multifactorial ANOVA was used to compare treatments (p-values 0.05 (*), 0.01 (**), 0.001(***) and 0.0001(****)).

### Chitosan induces *de novo* exudation of SA and phenolics in roots

In view of the results obtained with tomato roots elicited for 3 days with chitosan, we irrigated tomato plants with a low dose of chitosan (0.1 mg·mL^-1^) during 10, 20 and 30 days and analysed *de novo* exudation in roots. Ten days after planting (dap), chitosan increases *de novo* (p<0.01) exudation of a fluorescence signature putatively belonging to SA (Figure 4, Table 1). Twenty dap, chitosan significantly increases (p<0.001) fluorescence intensity of Component 3, putatively assigned to phenolics (Mostofa *et al.*, 2013; Parri *et al.*, 2020) and SA derivatives (Street and Schenk, 1981). This tendency is also found for Components 1 (putatively aromatic amino acids and peptides) and 2 (putatively IAA; Li *et al.*, 2009), although differences are not significant. In late root exudates (30 dap), fluorescence intensity decreases respect to early root exudates. This could be due to root ageing and lignification. At this time, EEM Fluorescence spectrum is separated into 2 Components only (Supplemental Table 1, Supplemental Figure 2) with no differences respect to controls.

**Figure 4.**
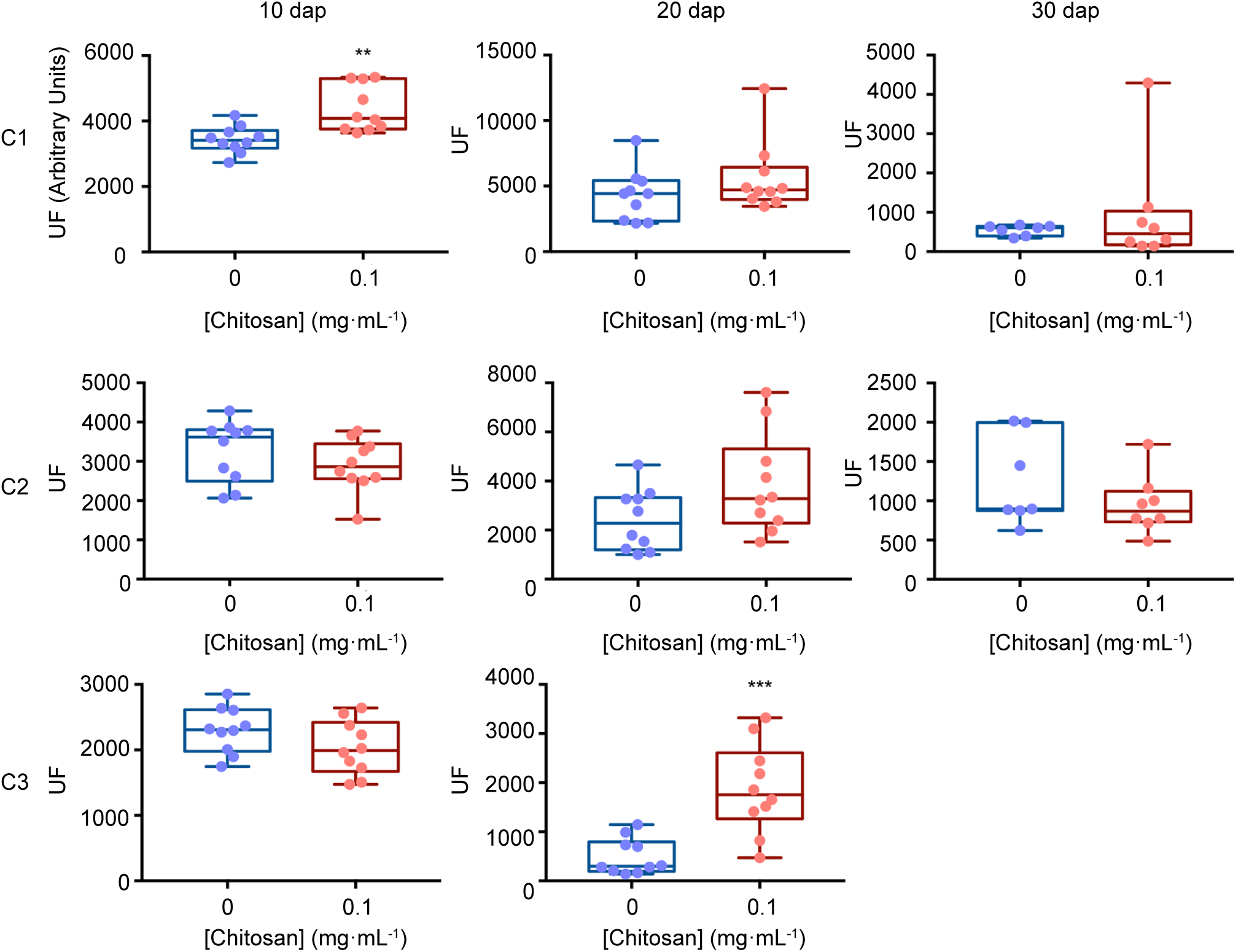
EEM fluorescence dynamics of tomato root exudates. For Ex/Em coordinates of Components see Table 1. Abbreviations: EEM = Emission Excitation Matrix, dap = days after planting, C1 = Component 1, C2 = Component 2, C3 = Component 3, UF = Fluorescence Units. Mann-Whitney test was used for not normal distribution treatments. Welch’s parametric test was used for the remaining treatments (p-values 0.05 (*), 0.01 (**), 0.001(***) and 0.0001(****)).

### Metabolomic diversity of tomato root exudates varies with time

Pools were made to concentrate root exudates and avoid sample variability (Yuan *et al.*, 2015; Yang *et al.*, 2016). Root exudates from 20 dap tomato plants (chitosan treated and controls) displayed most NMR peaks (Figure 5). This may be because older plants (30 dap) are more lignified (Cervilla *et al*., 2009) and display less rhizodeposition. Manual curation of ^1^H NMR profiles of root exudates from plants 10, 20 and 30 dap from chitosan treated plants and controls show no qualitative differences and contain 31, 123 and 18 peaks respectively. Ninety-three peaks are specific to 20 dap root exudates. Nine 20 dap specific peaks were identified as (Table 2): leucine/isoleucine (13), acetate (53), raffinose (99, 103), glucose (100), uracil (105 and 120), cinnamic acid (109, 119, 121), fumaric acid (109), p-aminobenzoic acid (112 and 122) and trigonelline (125, 130, 131). Chitosan decreases (p<0.05) acetate (peak 53) content in 20 dap root exudates. Six peaks, among them methanol (80) and formic acid (129), are found 20 and 30 dap. Lactate (peak 36) and malate (peaks 62 and 64) have been detected in all time points evaluated. All detected peaks are listed in Supplemental Table 2. A representative ^1^H NMR profile of tomato root exudates is shown in Supplemental Figure 3.

**Table 2.** Peak assignments for ^1^H NMR spectra of tomato root exudates. This table includes identified peaks only. Spectra part I, II and III are indicated in supplemental figure 3. Abbreviations: dap = days after planting, H mult. = H multiplicity, d = doublet, s = singlet, t = triplet, dt = double of triplets, m = multiplet. Treatment abbreviations: 0 = Tomato Root Exudates Control, 0.1 = Tomato Root Exudates from plants irrigated with 0.1 mg·mL^-1^ chitosan. Numbers in treatments correspond to the mean of maximum intensity of the peaks (arbitrary units). Bold numbers indicate significant differences (ANOVA, p<0.05).

| Part of the spectra | dap | Peak number | Compound | Shift (ppm) | H mult. | 20 dap [Chitosan]<br>($\text{mg}\cdot\text{mL}^{-1}$) | |
| --- | --- | --- | --- | --- | --- | --- | --- |
|  |  |  |  |  |  | 0 | 0.1 |
| I | 20 | 13 | Leucine/Isoleucine | 0.94 |  | 0.25475 | 0.17575 |
| I | 10-20-30 | 36 | Lactate | 1.33 | d | 0.35685 | 0.18225 |
| I | 20 | 53 | Acetate | 1.92 | s | 4.7895 | **3.4135** |
| I | 10-20-30 | 62 | Malate | 2.3 | t | 0.6217 | 0.5769 |
| I | 10-20-30 | 64 | Malate | 2.41 |  | 1.725 | 1.26 |
| II | 20-30 | 80 | Methanol | 3.36 | s | 0.34845 | 0.2379 |
| II | 20 | 99 | Raffinose | 4.99 | d | 0.027535 | 0.03025 |
| II | 20 | 100 | Glucose | 5.237 | d | 0.02523 | 0.035445 |
| II | 20 | 103 | Raffinose | 5.42 | d | 0.02462 | 0.02018 |
| II | 20 | 105 | Uracil | 5.79 | d | 0.031275 | 0.02049 |
| III | 20 | 109 | Cinnamic Acid / Fumaric Acid | 6.52 / 6.51 | d / s | 0.2005 | 0.15425 |
| III | 20 | 112 | p-Aminobenzoic Acid | 6.82 | d | 0.0033845 | 0.0050605 |
| III | 20 | 119 | Cinnamic Acid | 7.43 | m | 0.049535 | 0.04216 |
| III | 20 | 120 | Uracil | 7.52 | d | 0.00791 | 0.016235 |
| III | 20 | 121 | Cinnamic Acid | 7.62 | dt | 0 | 0.002784 |
| III | 20 | 122 | p-Aminobenzoic Acid | 7.73 | d | 0.0056825 | 0.00603 |
| III | 20 | 125 | Trigonelline | 8.07 | m | 0.006835 | 0.01239 |
| III | 20-30 | 129 | Formic Acid | 8.44 | m | 0.35505 | 0.1718 |
| III | 20 | 130 | Trigonelline | 8.82 | m | 0.028845 | 0.022265 |
| III | 20 | 131 | Trigonelline | 9.13 | s | 0.03144 | 0.024555 |

**Figure 5.**
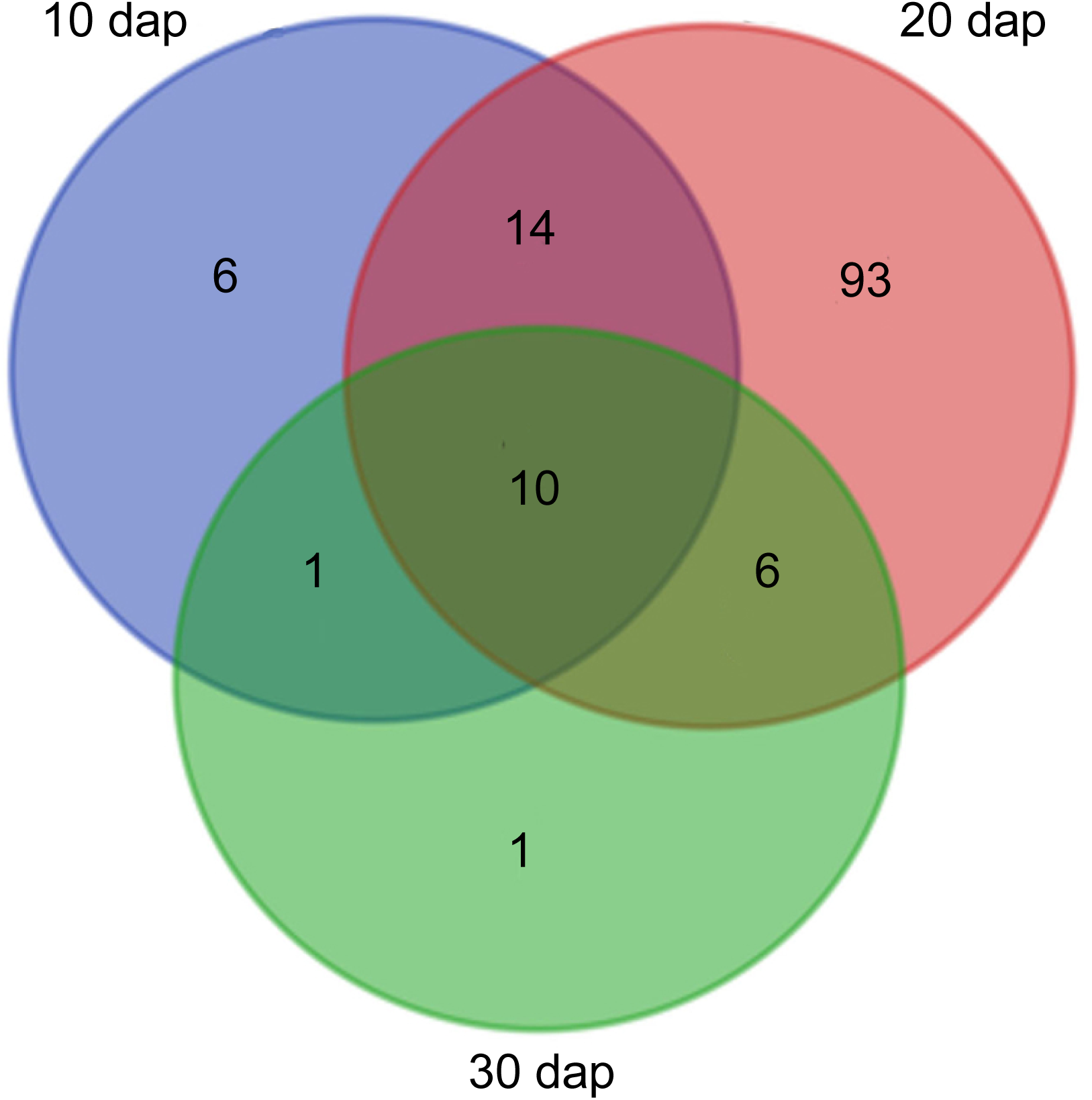
Tomato rhizodeposition varies with time. Venn-diagram of 1H Nuclear Magnetic Resonance analyses peaks detected in tomato root exudates at 10, 20 and 30 days after planting.

### Chitosan induces lipid signalling and defence compounds in tomato root exudates

Lipid signalling and defence compounds are putatively identified in 20 dap root exudates (Figure 6). Oxylipins and other lipid signalling compounds, such as 13-epi-12 oxophytodienoic acid (OPDA, 293.7, +), myo-inositol-2,4-diphosphate (IDP, 341.7, +) are up-regulated with chitosan. Two unidentified fatty acids (FA1, 313.8, +; FA2, 344.3, +) are slightly changed. Defence compounds such as glycerol-3-phosphate (G3P, m/z 191.0, mode -) and indol-3-carboxylic acid (I3CA, 290.8, +), involved in plant immunity are up-regulated with chitosan. On the contrary, 3-(2 methyl propyl) pyridine (MPP, 282.7, +), is down-regulated. Glucaric acid (GA, 327.7, -), which interacts with the pentose pathway, is also down-regulated.

**Figure 6.**
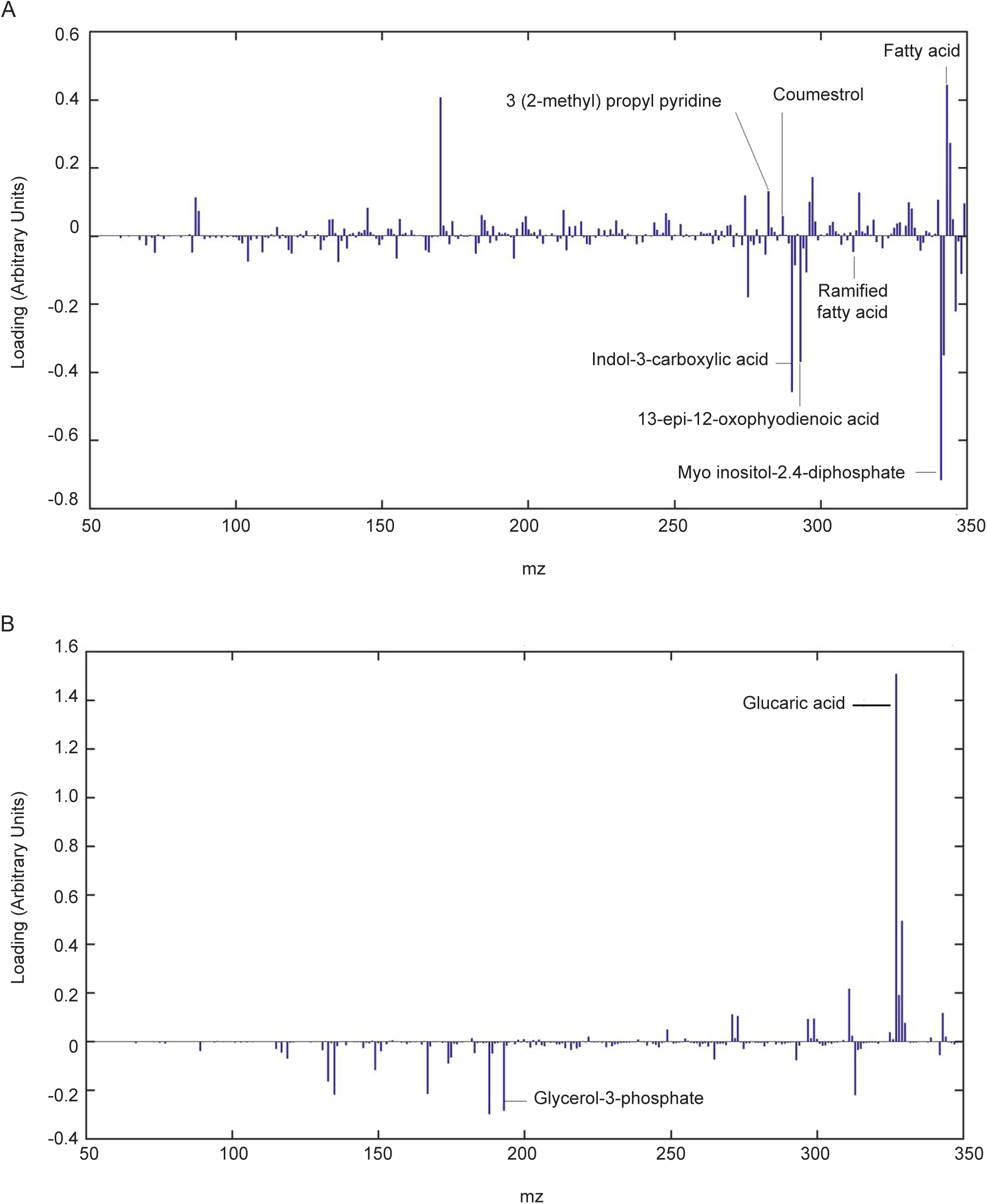
Chitosan induces lipid signalling and defence compounds in tomato root exudates. Partial Least Squares Regression Discriminant Analysis (PLSLDA) of HPLC-ESI-MS analysis of pools of 20 days after planting tomato root exudates in modes positive (A) and negative (B). Blue bars indicate a particular mass that differs from the control treatment. The larger the bar size, the more noticeable the difference in intensity of the mass compared to the control.

### Root exudates from plants treated with chitosan inhibit soil-borne pathogens

Soil-borne pathogens (fungi and nematodes) are inhibited by root exudates from plants treated with chitosan. Root exudates from chitosan-treated plants cause ca. 1.5-fold reduction (p<0.05) on hatching of the root-knot nematode *Meloydogine javanica* eggs after 72 hours respect to tomato control root exudates (Figure 7A). These exudates also inhibit growth of *Fusarium oxysporum* f.sp. radicis-lycopersici (FORL) ca. 2-fold respect to controls (Figure 7B). The chitosan resistant fungus *Pochonia chlamydosporia* strain 123 (Pc) does not show significant differences in growth with both exudates over time (Figure 7C).

**Figure 7.**
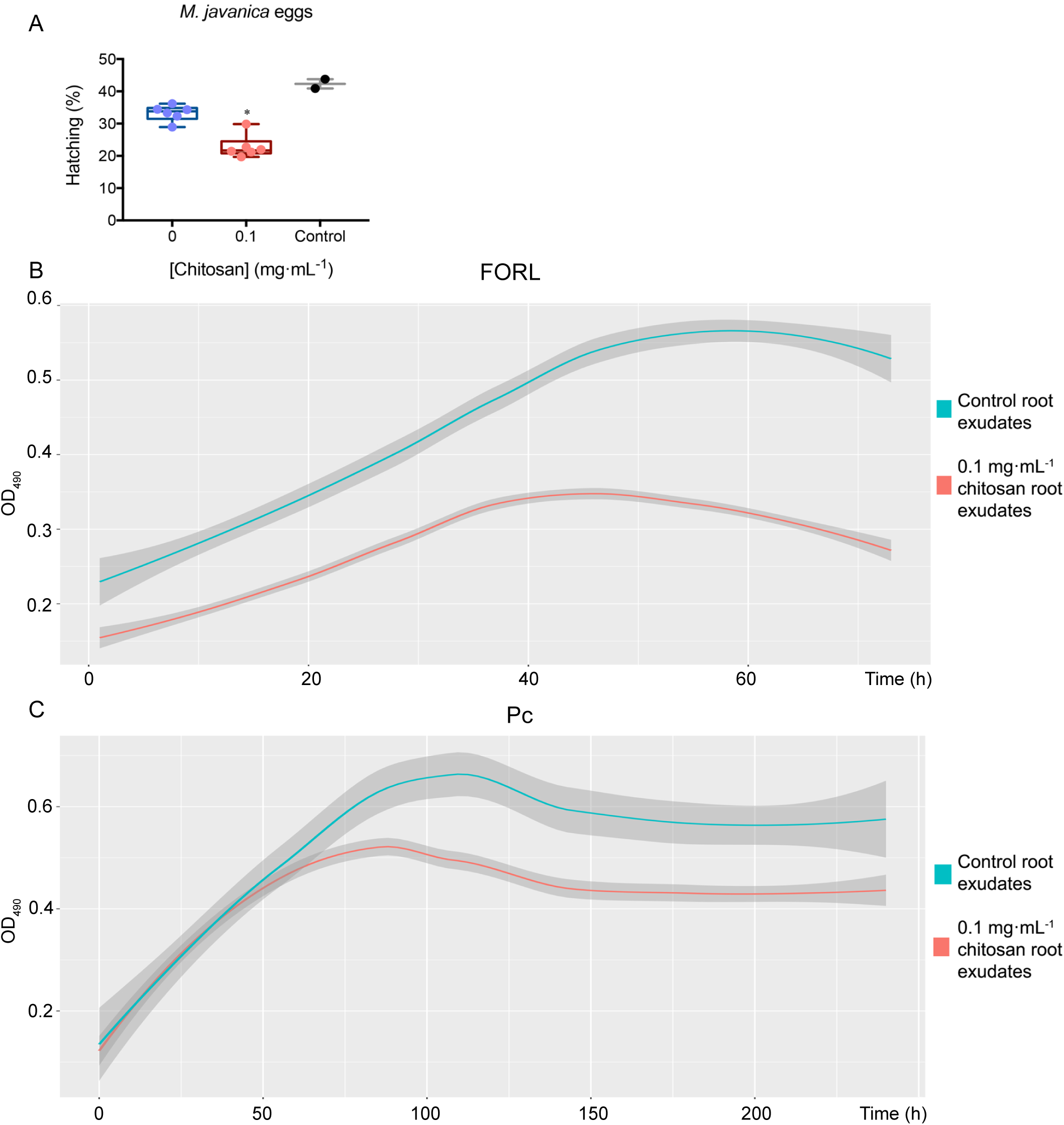
Root exudates from plants treated with chitosan inhibit soil-borne pathogens. A, Effect on hatching of Meloidogyne javanica eggs in tomato root exudates. Egg hatching is reduced ca. 1.5-fold. Multifactorial analysis Kruskal-Walis was performed to compare treatments (p-values 0.05 (*), 0.01 (**), 0.001(***) and 0.0001(****)). B, Growth kinetics of FORL in root exudates, chitosan inhibits ca. 2-fold growth. C, Growth kinetics of P. chlamydosporia strain 123 in tomato root exudates. Smoother Models were adjusted with band 0.75 and degree 2. Shadow area stablishes the prediction confidence intervals at 95%.

## DISCUSSION

In this study we give evidences that chitosan induces plant defences in tomato root exudates. High doses of chitosan are known to affect root growth (Lopez-Moya *et al.*, 2017; Asgari-Targhi *et al.*, 2018). Plasma membranes are likely to be a main target of this molecule (Palma-Guerrero *et al.*, 2010; Meisrimler *et al*., 2011; Jaime *et al.*, 2012). In this work, we show that increasing chitosan concentrations depolarize tomato root cells plasma membrane and reduce viability. He *et al*., (2009) propose chitosan as a potential cell penetration enhancer. This fact has been demonstrated using vital staining in fungi (Palma-Guerrero *et al.*, 2010) and, in this work, in root apices. These effects of chitosan on roots are likely to modify rhizodeposition (Pitta-Alvarez and Giulietti, 1999). Chitosan depolarisation on the membrane could trigger an increase of reactive oxygen species (ROS) in the plant cell, resulting in a signal cascade that modifies metabolic response (Pandey, 2017). ROS are known to alter chemical components of the cell and lipids in particular. Consequently, lipid signalling compounds, including oxylipins (OPDA and IDP) are up-regulated in root exudates from plants treated with chitosan (Dar *et al.*, 2015). OPDA, a precursor of JA (Dave and Graham, 2012), could explain the up-regulation of JA in these exudates. OPDA is involved in plant defense by up-regulating detoxifying enzymes and secondary metabolism pathways (Conconi *et al*., 1996; Savchenko *et al*., 2014). Oxylipins other than jasmonates are probably also essential for the resistance of plants to pathogens (Blée, 2002). IDP, which belongs to a family of compounds that act as messengers that regulate cellular functions, including cell cycling, apoptosis, differentiation, and motility (Majerus, 1992) is also up-regulated by chitosan. IDP is well-known intermediate of metabolic and signalling pathways (Berridge, 1993;Williams *et al*., 2015; Freed *et al*., 2020). Besides IDP, other lipids (eg. ramifed fatty acids) found in root exudates, could be part of the plant immune response signalling induced by chitosan (Czékus *et al*., 2020; Narula *et al*., 2020). Other plant defence compounds such as G3P and I3CA are also up-regulated with chitosan. G3P is systemic acquired resistance (SAR) inducer in plants (Chanda *et al*., 2011; Wang *et al*., 2019). I3CA, an indolic compound, also induces plant resistance (Böttcher *et al*., 2014; Syeda and Riazunnisa, 2020) through callose accumulation (Pastor-Fernández *et al*., 2019). On the other hand, GA, which interacts with the pentose pathway (Moon *et al*., 2009), is down-regulated by chitosan. We could assume that chitosan is causing plant stress and that interferes with its primary metabolism and reduces pentose phosphate pathway.

Importance of chitosan in plant hormone production and systemic acquired response has been widely demonstrated (Mika *et al*., 2010; Iglesias *et al*., 2019; Colman *et al*., 2019; Fooladi-Vanda *et al*., 2019; Ma *et al*., 2019). We have found that increasing chitosan leads to the accumulation of fluorescence compounds corresponding to phenolics, as well as hormones related to plant growth and defence (JA, SA, ABA and IAA). This correlation between chitosan and phenolics has been previously studied (Pitta-Alvarez and Giulietti, 1999; Park *et al*., 2019; Jaisi and Panichayupakaranant, 2020; Samari *et al*., 2020). Chitosan enhances metabolic pathways (e.g. phenylpropanoid) involved in the biosynthesis of phenolic compounds (Asgari-Targhi *et al*., 2018; Fooladi-Vanda *et al*., 2019; Singh *et al*., 2020). A low dose of chitosan enhances the plant immune response by plant hormone (JA and SA, mainly) accumulation in root tissues (Lopez-Moya *et al*., 2017; Iglesias *et al*., 2019; Singh *et al*., 2020). However, this effect has not yet been detected in tomato root exudates. In addition, there is evidence that chitosan could be used as a substitute for commonly used growth factors such as methyl jasmonate, auxins or cytokinins (Cui *et al*., 2012; Sivanandhan *et al*., 2012; Ahmad *et al*., 2019; Acemi, 2020) due to its elicitor effects (Salachna and Zawadzińska, 2014; Malerba and Cerana, 2016). Phytomelatonin has been recently considered a plant master regulator involved in abiotic and biotic responses. In our work, exposure of tomato roots to increasing chitosan doses accumulates phytomelatonin and hormones in roots and root exudates in a typical biotic stress response (Arnao and Hernández-Ruiz, 2019c; Moustafa-Farag *et al.*, 2019). The decrease in levels of some plant hormones at toxic chitosan doses has also been found for phenolics (Park *et al.*, 2019). EEM Fluorescence signatures corresponding to IAA (Li *et al.*, 2009) and SA (Miles and Schenk, 1970) are increased by chitosan in tomato root exudates. These hormones are known to be up-regulated by chitosan (Lopez-Moya *et al.*, 2017; Fooladi-Vanda *et al*., 2019). SA and JA are related to plant responses to stress (War *et al.*, 2011). Therefore, chitosan can be an elicitor of plant defences in root exudates (Benhamou, 1996). Our bioassays show that chitosan induces root exudates inhibitory to root pathogenic fungus FORL and root-knot nematode eggs without significantly affecting the growth and development of a biocontrol fungus (Pc). The toxic effect of chitosan-derived exudates may be related to the overproduction of SA which, in combination with chitosan induce SAR and reduce infection by root-knot nematodes (Vasyukova *et al*., 2003; Singh *et al*., 2019). Chitosan, moreover, by itself, is toxic to fungi such as *Fusarium spp.* (Palma-Guerrero *et al.*, 2010; Al-Hetar *et al*., 2011), and other studies show that plants treated with chitosan reduce FORL infection (Benhamou and Thériault, 1992).

Chitosan depolarizes plasma membrane of tomato root cells, causing the secretion of hormones, lipid signalling and plant defence compounds, including phenolics. This process affects cell viability and rhizodeposition, so plant age must be considered before applying chitosan, as well as the duration of applications. Our results have proven that root exudates of chitosan-treated plants are able to reduce soil-borne pathogens growth and development, which makes chitosan a promising tool for preventing and managing pests and diseases in a sustainable way.

## MATERIALS AND METHODS

### Chitosan, plants, fungi and nematodes

Chitosan with 70 kDa molecular weight and 80.5% deacetylation degree was used in all experiments. Chitosan was obtained from Marine BioProducts GmbH (Bremerhaven, Germany) and made as in Palma-Guerrero *et al*. (2007). Tomato plants (*Solanum lycopersicum* cv. Marglobe) were used in all experiments. *Pochonia chlamydosporia*, isolate number 123 (ATCC no. MYA-4875; CECT no. 20929) was isolated from *Heterodera avenae* eggs in south west Spain (Olivares-Bernabeu and Lopez-Llorca, 2002). *Fusarium oxysporum* f. sp. radicis-lycopersici (CBS 123668) was obtained from CBS-KNAW culture collection. *Meloidogyne javanica* was obtained from a field population and maintained in susceptible tomato plants. Nematode egg masses were dissected from root-knot nematode infested roots and stored at 4°C. Egg masses were hand-picked and surface-sterilized as in McClure *et al.* (1973) with modifications.

### Root electrophysiology experiments

Tomato seeds were surface sterilized using 1% sodium hypochlorite and washed three times 1 min each with sterile distilled water (SDW). Tomato seeds were then placed on germination medium (GM, Glucose 10 g·L^-1^, Yeast Extract 0.1 g·L^.1^, Bactopeptone 0.1 g·L^-1^, Technical Agar 12 g·L^-1^). GM plates with tomato seeds were placed at 4°C for 2 days for stratification and incubated at 24°C, 65% relative humidity (RH), in the dark for 5 days and in a photoperiod (16:8) for further 5 days. Plantlets were then placed individually in a holder chamber filled with Gamborgs’s B5 1:10 (Sigma; Gamborg *et al.*, 1968). Glass microcapillaries filled with 0.5 M KCl (tip diameter < 1 µm) were inserted into root cortical cells until a stable basal potential was obtained (Gunsé *et al*., 2016). Roots were exposed to increasing concentrations of chitosan (0.1, 1 and 2 mg·mL^-1^ in Gamborgs’s B5 1:10 medium) and membrane potentials were recorded. No chitosan was used for control treatments. Between each change of chitosan concentration, root medium was replaced for Gamborgs’s B5 1:10 to check the physiological status of the cell. At least 3 replicates were performed per treatment.

### Root vital staining

Tomato plantlets were placed in Gamborgs’s B5 1:10 liquid medium for one day for acclimatization. Chitosan then was added at 0.1, 1 and 2 mg·mL^-1^. Plants exposed only to Gamborg’s B5 1:10 were used as controls. Plantlets with treatments were incubated for 24 h. Roots were then stained with fluorescein diacetate (FDA), 5 mg·mL^-1^ in acetone diluted 1:250 in Dulbecco’s phosphate-buffered saline (DPBS, ThermoFisher) and propidium iodide (PI, 20 mg·mL^-1^ in DPBS; Jones and Senft, 1985). Roots were visualized using a Nikon Optiphot microscope using 10X objective with an attached Nikon DS–5 M camera system using epifluorescence (Ex: 450-490 nm, Em: 520 nm). Non-damaged cells show green FDA fluorescence and nuclei of damaged cells show red PI fluorescence.

### Plant growth conditions

Tomato plantlets (Experiment 1) were placed in 200 mL expanded polystyrene sterile cups each containing 100 cm^3^ of sterilized sand. They were then incubated in a culture chamber (SANYO MLR-351H) at 65% RH, 24°C with a 16:8 h (light:dark) photoperiod. Plants were irrigated for 20 days with Gamborg’s B5 basal mixture 1:10 keeping moisture to field capacity. Plants were then removed, washed and introduced individually in Magenta Boxes (575 mL, Sigma). Each Box was filled with 50 mL of Gamborg’s B5 1:10 (control) or Gamborg’s B5 1:10 amended with chitosan (0.1, 1, 2 mg·mL^-1^). Nine tomato plants were set per treatment. Plants were incubated for 3 days as described above.

In Experiment 2, tomato plantlets were grown as for Experiment 1. Plants in cups were then irrigated with 1:10 Gamborg’s B5 basal mixture on its own or amended with 0.1 mg·mL^-1^ chitosan. Plants were incubated as before. Ten plants per treatment were sampled 10, 20 and 30 days after planting (dap) for further analyses.

### Collection of root exudates

In Experiment 1, root exudates accumulated for 3 days in Magenta Boxes were collected and filtered through Miracloth (Calbiochem). Pools were made with the exudates of 3 plants each. Root exudate pools were frozen at −20°C until further use. In Experiment 2, whole plants were removed, and their root systems washed in SDW. *De novo* root exudates from these plants were collected by placing individual whole plants in sterile plastic containers with 20 mL SDW per gram of root. Plants were incubated in the dark at 24°C, 65% RH for 24 h. Plants were then removed, and root exudates were collected by 0.22 μm (Q-MAX) filtration and stored frozen at −20°C until used.

### Emission Excitation Matrix (EEM) Fluorescence analysis

Two mL of each root exudate (Experiments 1 and 2) were collected and EEM Fluorescence spectra were obtained with a spectrofluorometer (Jasco FP-6500) equipped with a 150W Xenon lamp. Contour maps of EEM fluorescence spectra were obtained from water extracts of whole root exudates (10 samples) or pools. The emission (Em) wavelength range was fixed from 220 to 460 nm in 5 nm steps, whereas the excitation (Ex) wavelength was fixed from 220 to 350 nm in 2 nm steps. The slit width was 5 nm and the root exudates were placed in a 1cm path length fused quartz cell (Hellma). The UV-visible spectra of samples were acquired (SHIMADZU UV-160 spectrophotometer, 200-800 nm, 1cm quartz cuvette). Absorbance was always lower than 0.1 (OD_units_) at 254 nm in order to reduce the absorbance of the solution to eliminate potential inner filter effects (Mobed *et al.*, 1996). EEM fluorescence spectra of root exudates were analysed using Parallel Factor Analysis (PARAFAC) as in Ohno and Bro (2006). PARAFAC model Components were calculated for each treatment and time.

### Plant hormone analysis

Root exudate pools (Experiment 1) were lyophilized to analyse Indoleacetic Acid (IAA), Abscisic Acid (ABA), Salicylic Acid (SA) and Jasmonic Acid (JA) by Ultra Performance Liquid Chromatography-Mass Spectrometry (UPLC-MS). Material was extracted with 80% Methanol-1% Acetic acid. Deuterium-labelled hormones (purchased from Prof. L Mander-Canberra, OlChemim Ltd. -Olomouc): [^2^H_5_] IAA, [^2^H_4_] SA and [^2^H_6_] ABA were added as internal standards. For quantification of JA, dhJA was used instead. For collecting the fractions containing SA, ABA and JA; extracts were passed consecutively through HLB (reverse phase), MCX (cationic exchange) and WAX (ionic exchange) columns (Oasis 30 mg, Waters), as described in Seo *et al.* (2011). The final residue was dissolved in 5% Acetonitrile - 1% Acetic acid and separated by reverse phase UPHL chromatography (2.6 µm Accucore RP-MS column, 100 mm length x 2.1 mm i.d.; ThermoFisher Scientific) with a 5% to 50% acetonitrile gradient. Hormones were analysed by electrospray ionization and targeted-SIM using a Q-Exactive spectrometer (Orbitrap detector, ThermoFisher Scientific). Concentrations of hormones in extracts were determined using embedded calibration curves and the Xcalibur 4.1 SP1 build 48 and TraceFinder programs. We thank Dr Esther Carrera by hormone quantification carried out at the Plant Hormone Quantification Service, Valencia, Spain (www.ibmcp.upv.es).

Phytomelatonin was also tested in both exudates and tissues. Ten-day old seedlings were grown with chitosan (0.1, 1, 2 mg·mL^-1^) in Gamborg’s B5 1:10 for 3 days. Plants exposed to Gamborg’s B5 1:10 only were used as control. For each treatment, 0.2-0.3 g of roots were dried onto sterile paper and placed in a 4 mL polypropylene tube with 3 mL ethyl acetate. Samples were shaken at 120 rpm and 4°C in the dark overnight. Roots were removed and solvent evaporated under vacuum. The dry residue was resuspended in 1 mL acetonitrile and filtered by 0.22 µm. Phytomelatonin from root tissues was quantified by HPLC with fluoresecence detection with Ex/Em wavelength pair of 280/348 nm, as in Hernández-Ruiz *et al*. (2005). For root exudates, three samples of 1 mL of Experiment 1 pools were used. Phytomelatonin was extracted and analysed as described above.

### Nuclear Magnetic Resonance (^1^H NMR)

Root exudates from Experiment 2 were pooled 5 by 5, lyophilized and resuspended in 1 mL of D_2_O (deuterated water). Two pools were obtained from each treatment and time.

Six hundred microliters of filtered pools were placed in a 5 mm NMR tube with 0.75% 3-(trimethylsilyl)propionic-2,2,3,3-d_4_ acid sodium salt (TSP) and 0.002 g sodium azide. ^1^H NMR experiments were performed on a Bruker AVIII 700 MHz (CITIUS, University of Sevilla, Spain). The number of scans was 256 and the experiments were carried out at 298 K. ^1^H chemical shifts were internally referenced to the TSP at δ 0.00. ^1^H NMR spectra were aligned using TopSpin™ (Bruker).

Sensitivity of NMR is different in each region. For region I (organic-acid and amino-acid region), the threshold was set in 0.1. For II (sugars/polyalcohols region) and III (phenolics/aromatic compounds region) peaks showed less intensity, consequently the threshold was set in 0.01 and 0.001 respectively.

### High Performance Liquid Chromatography Electrospray Ionization tandem Mass Spectrometry (HPLC-ESI-MS)

HPLC-ESI-MS analyses were performed with a High-Performance Liquid Chromatography system with an Agilent 1100 Series model coupled to a UV-visible variable wavelength detector and a mass spectrometer with ion trap analyser Series LC/MSD Trap SL (Agilent, Santa Clara, CA). The mass spectrometer was operated in the positive and negative ESI modes, and the ion spray voltage was set at 4 kV. Mass range was set from 50 to 350 atomic mass units. Nitrogen was used as carrier gas (70 psi), and the ion transfer capillary heated to 350°C. Injections were carried out using an HTC Pal autosampler (CTC Analytics, Zwingen, Switzerland) equipped with a 20 μL sample loop.

Pools of 20 dap tomato root exudates (Experiment 2) were infused into the flow of the HPLC system (10 μL) through a T connection under the following conditions: flow rate, 1 mL·min^-1^. Ultrapure water with 0.1% Formic Acid was used as Solvent A, whereas MeOH with 0.1% Formic Acid was used as Solvent B. From 0-15 min Solvent B was kept at 10% and afterwards a gradient to 90% was established during 20 min and decreased again to 10% until minute 25. The LC separations were carried out with a Poroshell 120 EC-C18 column, (4.6 x 100mm, 2.7μm -Agilent Technologies-). Runs were performed at 25 °C. Raw data were transformed as explained in Marhuenda-Egea *et al.* (2013).

Partial Least Squares Regression Discriminant Analyses (PLSLDA) show that m/z intensities depend on treatment. Loading values lower than 0 indicate that m/z intensity is higher in chitosan-treated plant root exudates. A threshold was set at ±0.2 to identify m/z with significant variations. Datasets were placed in Dryad repository (https://doi.org/10.5061/dryad.ghx3ffbkb).

### Evaluation of tomato root exudates on fungi and nematode eggs

Bioassays were performed in 96-well plates (Thermo Scientific) using 20 dap root exudate pools obtained in Experiment 2. For fungal experiments, 200 μL of exudate and conidia were added per well to reach a concentration of 10^6^ conidia·mL^-1^. Each exudate was analysed by triplicate and for each treatment, there were two pools of exudates. For both, FORL and Pc, OD_490_ was calculated after 4 and 8-days respectively using a microplate reader (Tecan SPECTRAfluor). Results were handled with XFluor Software™. For time series with FORL and Pc, the relation between two variables is not linear. In both cases, Smoother Model Lowess (“Locally weighted regression”) was applied (Cleveland, 1979).

To evaluate the effects of root exudates on root-knot nematode eggs, experiments were performed with 200 μl of exudate or water (control) containing 100 *M. javanica* eggs each. Hatching percentage was scored using an inverted microscope after 72 h incubation at 30°C.

### Statistical analyses

EEM Fluorescence data was analysed by PARAFAC, as described above, and the contribution of the Components 1, 2 and 3 analysed by ANOVA tests. The level of significance in all cases was 95%. All statistical analyses were performed using GraphPad Prism version 7.00 (GraphPad Software, La Jolla California, USA, https://www.graphpad.com/).

HPLC-ESI-MS data were processed using a Partial Least Square (PLS) regression model (Verboven and Hubert, 2005; Marhuenda-Egea *et al.*, 2013) Classical PCA were also performed to display and group data (Verboven and Hubert, 2005). This data analysis was carried out using the LIBRA toolbox (available at http://wis.kuleuven.be/stat/robust/software).

## Acknowledgements

This work was supported by AGL 2015 66833-R Grant from the Spanish Ministry of Economy and Competitiveness and H2020 MUSA 727624 European Project. Thanks are due to members of the Plant Pathology Laboratory of the University of Alicante, for their help and support. Authors also wish to thank Mr Arnau Hernández Royo for his help conducting electrophysiological and microscopy experiments.

